# Plasma-Activated Water: An Emerging Technology for the Control of Eggshell-Associated Biofilms

**DOI:** 10.64898/2026.01.12.699152

**Authors:** Heema Kumari Nilesh Vyas, Adrian Issa Abdo, Bjoern Hendrik Kolbe, Angela Boahen, Siyuan Jia, Andrea Rene McWhorter, Bryan Robert Coad, Katharina Richter

## Abstract

Biofilm formation on eggshell surfaces poses a significant food safety concern by promoting bacterial persistence and facilitating cross-contamination, contributing to foodborne illness outbreaks worldwide. Beyond public health impacts, bacterial contamination also drives spoilage, food loss, and waste, together placing substantial strain on healthcare systems and economic infrastructures. This study evaluated the anti-biofilm efficacy of plasma-activated water (PAW), a reactive oxygen and nitrogen species (RONS)–rich solution generated through the interaction of cold atmospheric plasma with water. 50 bacterial isolates were recovered from chicken eggshells, and the biofilm-forming capacities of eight representative species (*Staphylococcus arlettae, Lysinibacillus fusiformis, Priestia megaterium, Peribacillus frigoritolerans, Acinetobacter lwoffii, Escherichia coli, Pseudomonas fulva,* and *Pseudomonas stutzeri*) were characterised *in vitro*. All isolates formed biofilms, exhibiting variable culturable cell populations, biomass, and extracellular polymeric substance (EPS) matrix presence and composition. EPS analyses identified polysaccharides, extracellular DNA, and proteins as key matrix components, with relative abundance varying with biofilm maturity. PAW efficacy was assessed under short exposure times (1, 5, and 15 minutes) using freshly generated, 1-week-old, and 4-week-old PAW. Fresh PAW demonstrated potent anti-biofilm activity, achieving complete inactivation of *E. coli* and *P. stutzeri* biofilms below the limit of detection within 1 min, while biofilms of Gram-positive bacteria, *L. fusiformis* and *P. megaterium*, were generally more tolerant. Reduced efficacy of aged PAW was likely associated with RONS decay. Mechanistic analyses, including scanning electron microscopy and intracellular reactive oxygen species assays, linked PAW treatment to biofilm disruption and intracellular oxidative stress. This study provides the first detailed evaluation of biofilm formation by under-investigated eggshell-associated bacteria and elucidates mechanisms underlying PAW-mediated biofilm inactivation. Overall, PAW shows strong potential as a sanitising technology for the egg industry, supporting safer food production and reduced reliance on conventional antimicrobial agents amid rising global antimicrobial resistance rates.

## 1. Introduction

Fresh eggs and egg-derived products represent a major component of the global poultry industry. China is recognised as the world’s largest egg producer and consumers, accounting for approximately one-third of global egg production^1^. In India, annual egg production is estimated at 74 billion eggs^2^, while the United States produces approximately 110 billion eggs^3^. During the 2024–25 financial year, Australia’s egg production was estimated to be around 7 billion eggs^4^.

Although eggs are generally classified as a medium-to-low risk food for foodborne illness, egg-related outbreaks continue to contribute substantially to public health and economic burdens, accounting for an estimated US$2 billion in annual foodborne disease–related costs in Australia alone^5^. This moderate risk classification primarily reflects the prevalence of *Salmonella* (largely *Salmonella enterica* serovars Enteritidis and Typhimurium) as the leading cause of egg-associated food poisoning. Globally, *Salmonella* infections are estimated to cause tens of millions of illnesses and around 230,000 deaths annually, many of which are linked to foodborne transmission^6, 7^. In addition to foodborne illness, bacterial contamination of eggshell surfaces contributes to spoilage and reduced shelf-life, increasing food loss and waste across the egg supply chain. Other members of the Enterobacteriaceae family, including *Escherichia coli* and other coliforms, as well as non-Enterobacteriaceae genera such as *Pseudomonas*, *Acinetobacter*, *Staphylococcus*, and *Bacillus*, are frequently isolated from eggshells and processing environments and are associated with increased health safety risks and quality deterioration^3^.

Egg contamination occurs via both vertical and horizontal transmission routes. Vertical transmission occurs when certain pathogens, particularly specific *Salmonella* serotypes, colonise the oviduct of laying hens, leading to internal egg contamination prior to shell formation. Conversely, horizontal transmission occurs post-lay, when eggs come into contact with contaminated materials such as faeces, dust, soil, water, nesting material, or other broken eggs^8^, facilitating bacterial adherence and colonisation of the shell surface. Despite widespread on-farm control strategies, downstream contamination during grading and packing remains a persistent challenge. Contaminated equipment, regardless of good personnel hygiene practice, can enable bacterial transfer regardless of stringent sanitation protocols^9^. A large-scale survey of six poultry processing plants detected *Salmonella* on equipment after processing (36%), after cleaning (12%), and even after sanitation (9%), underscoring the risk of cross-contamination from surfaces to eggs and other poultry products^10^.

The persistence of *Salmonella* and other bacteria on eggshells has been increasingly attributed to biofilm formation^11–13^. Many foodborne pathogens, including *E. coli*, *Staphylococcus aureus*, and thermophilic *Campylobacter*, predominantly exist within aggregated microbial communities rather than as free, single (planktonic) cells^3, 14, 15^. Biofilms are complex, often polymicrobial structures comprising of microbial cells embedded in an extracellular polymeric substance (EPS) matrix composed of polysaccharides, extracellular DNA (eDNA), proteins, and lipids^16^. This matrix provides considerable protection against biological, chemical, and physical stressors encountered in food-processing environments. Moreover, biofilm-associated cells exhibit altered metabolism and gene expression, resulting in 10–1000-fold greater tolerance to antimicrobials than the planktonic form^17, 18^.

In commercial egg production, washing is an essential step to remove debris and reduce microbial load before packing. This rapid process (≈30 seconds) typically involves pre-washing with warm water, washing with surfactants, sanitising with antimicrobial agents (typically chlorine-based; ≈100–200 ppm), and blow-drying^19^. However, the porous structure of eggshells allows microbial contaminants and chemical sanitising solutions, such as chlorine, to permeate into the egg contents, raising additional concerns regarding food safety, chemical residues, and public health. Combined with the persistence of biofilms and the growing threat of antimicrobial resistance (AMR), these issues underscore the urgent need for safer and more effective sanitation technologies and agents.

Plasma-activated water (PAW) has emerged as a promising non-thermal sanitisation and decontamination technology, either as a stand-alone intervention or as an adjunct to conventional food-washing systems. PAW is produced when cold atmospheric plasma (CAP), interacts with water, generating a solution enriched with reactive oxygen and nitrogen species (RONS). These include short-lived species (e.g., hydroxyl radicals, atomic oxygen, superoxide anions, peroxynitrite) and relatively longer-lived species (e.g., hydrogen peroxide, ozone, nitrite, nitrate)^20^. Plasma activation also induces other physicochemical changes such as decreased pH, and increased conductivity and oxidation–reduction potential (ORP), all contributing to its antimicrobial activity. This complex composition underpins the potent, broad-spectrum antimicrobial properties of PAW, enabling rapid inactivation of bacteria as well as other microorganisms like viruses and fungi (including moulds, yeasts, and spores) through oxidative and nitrosative damage to cellular membranes, proteins, enzymes, and nucleic acids^21–32^. Because its effects are multi-targeted, PAW presents a low risk for AMR development.

The efficacy of PAW in reducing microbial loads on fresh produce, meat, poultry, and food-contact surfaces has been demonstrated in multiple studies against key foodborne pathogens such as *E. coli*, *Pseudomonas aeruginosa*, *Listeria monocytogenes*, *Salmonella* Typhimurium, and *S. aureus* biofilms^22, 24, 25, 33–35^. However, the ways in which PAW interacts with biofilms formed by eggshell-surface bacteria are still poorly understood. Compared with conventional sanitisers such as chlorine-based agents, quaternary ammonium compounds, peracetic acid, and hydrogen peroxide^25, 36, 37^, PAW offers several distinct advantages: it is generated solely from gas (e.g., air), water, and electricity; is purported to not produce harmful residues; and its physicochemical properties can be finetuned by adjusting plasma parameters (e.g., voltage, gas flow) or treatment modes (e.g., dipping, spraying, or rinsing). These features position PAW as a next-generation sanitising technology with potential to enhance or aid food safety, reduce harsh chemical use, and mitigate AMR within egg and broader food processing industries.

In this study, 50 bacterial isolates were recovered and identified from chicken eggshells. Eight total representative Gram-positive and Gram-negative species were characterised *in vitro* for their biofilm-forming capacities, including analysis of total culturable viable cells, biomass, and EPS presence and composition analysis (polysaccharides, eDNA, and proteins) across different biofilm ages. Biofilms grown for 48 h were used for antimicrobial susceptibility testing to PAW treatments of varying duration (1, 5, and 15 minutes) and age (fresh, 1-week-old, and 4-week-old PAW). Parallel physicochemical assessments of PAW enabled association of RONS profiles, pH, ORP, and conductivity with anti-biofilm efficacy. Mechanistic studies harnessing scanning electron microscopy for surface morphological changes and intracellular reactive oxygen species assays further elucidated PAW-induced biofilm structural and architectural changes as well as individual cellular effects.

## 2. Materials & Methods

### 2.1. Egg Collection & Isolation of Bacteria from the Eggshell Surface

Unwashed, intact, floor-laid chicken eggs (n = 12) were collected without washing or sanitisation from backyard laying hens maintained by four independent household keepers within the Adelaide metropolitan region (three eggs/household) and transported under ambient conditions for immediate processing. Individual eggs were placed separately into sterile polyethylene bags containing 10 mL of sterile buffered peptone water (BPW). Each egg was gently agitated in the BPW for 1 min at ambient temperature to recover eggshell-associated bacteria, after which the egg was aseptically removed and the rinsate retained for microbiological analysis. Specifically, 1 mL aliquots of each rinsate were plated in duplicate onto selective agar media as follows: Violet Red Bile Glucose agar (Thermo Scientific) and Brilliance™ *Salmonella* agar (Oxoid) for isolation of Enterobacteriaceae^38^; Brilliance™ CRE agar (Thermo Scientific) or CHROMagar™ *Acinetobacter* (CHROMagar) for *Acinetobacter* spp.^39^; and Pseudomonas Isolation Agar (Thermo Scientific™ Remel™) supplemented with glycerol for *Pseudomonas* spp^40^. Plates were incubated aerobically at 37 °C for 18–24 h.

From each positive sample, two presumptive colonies were randomly selected and purified by re-streaking on the corresponding selective medium. Presumptive identification was based on characteristic colony morphology: Enterobacteriaceae produced dark red–purple colonies with red–purple haloes (with *Serratia* spp. appearing blue on Brilliance™ *Salmonella* agar); *Acinetobacter* spp. appeared white or naturally pigmented on Brilliance™ CRE agar or red on CHROMagar™ *Acinetobacter*; and *Pseudomonas* spp. produced fluorescent green colonies with pyocyanin pigmentation. Purified isolates were streaked onto Heart Infusion (HI) agar and incubated overnight at 37 °C. Following incubation, isolates were preserved in glycerol stocks at −80 °C for long-term storage, while parallel agar-struck isolates were maintained at 4 °C for short-term storage prior to DNA extraction and preparation for subsequent sequencing.

### 2.2. Bacterial Isolate DNA Extraction & Sequencing

Genomic DNA was extracted from bacterial isolates recovered from eggshell surfaces using the Quick-DNA™ Fungal/Bacterial Miniprep Kit (Zymo Research, USA) according to the manufacturer’s instructions. Briefly, 50 purified isolates were randomly selected to represent the diversity of bacteria recovered from the eggs. Each isolate was cultured overnight in 10 mL HI broth at 37 °C with shaking at 160 rpm. Cultures were centrifuged at 7,000 × g for 10 min, and cell pellets were resuspended in ultrapure water and BashingBead™ buffer. Mechanical lysis was performed using a TissueLyser II (Qiagen) at 30 Hz for 5 min, repeated twice to ensure complete cell disruption. Following bead beating, samples were centrifuged at 10,000 × g for 1 min, and the supernatants were transferred to clean tubes, mixed with Genomic Lysis Buffer, and centrifuged again at 10,000 × g for 1 min. DNA purification was completed using the kit’s DNA Pre-Wash and Wash Buffers, and genomic DNA was eluted into clean collection tubes. DNA concentration and purity were assessed using a NanoDrop One spectrophotometer (Thermo Fisher Scientific), with yields (ng/µL) and A260/A280 ratios confirmed.

The 16S rRNA gene was amplified using universal primers 27F (5′-AGAGTTTGATCMTGGCTCAG-3′) and 1492R (5′-CGGTTACCTTGTTACGACTT-3′) with MyTaq™ Red Mix (Bioline). PCR amplification comprised an initial denaturation at 95 °C for 1 min, followed by 35 cycles of denaturation at 95 °C for 15 s, annealing at 64.3 °C for 15 s, and extension at 72 °C for 45 s, with a final extension at 72 °C for 10 min. A second PCR was performed using the first-round PCR product under identical cycling conditions. Nuclease-free water was included as a negative control, and genomic DNA from *Salmonella enterica* serovar Typhimurium ATCC 14028 was used as a positive control in each PCR run. PCR amplicons were excised from agarose gels and purified using the PureLink™ Quick Gel Extraction Kit (Invitrogen). Purified products were quantified using a NanoDrop™ One spectrophotometer and submitted to the Australian Genome Research Facility for Sanger sequencing. Resulting sequences were analysed using BLASTn (NCBI) and species-level identification was assigned based on BLAST scores and ≥99% sequence identity.

### 2.3. Biofilm Formation

From the 50 eggshell-associated bacterial isolates identified, a representative subset of four Gram-positive (*Staphylococcus arlettae, Lysinibacillus fusiformis, Priestia megaterium,* and *Peribacillus frigoritolerans*) and four Gram-negative (*Acinetobacter lwoffii, Escherichia coli, Pseudomonas fulva,* and *Pseudomonas stutzeri*) species was selected for further investigation. These species were selected to represent a combination of well-recognised foodborne or opportunistic pathogens and lesser-studied environmental bacteria commonly associated with soil, water, and poultry environments, for which biofilm formation remains poorly characterised. Single-species biofilms were developed in sterile, flat-bottom 96-well microtiter plates over 24, 48, and 72 h to assess formation dynamics and determine the optimal growth period. Following Vyas *et al* (2021)^41^, wells were inoculated with 150 µL of diluted overnight bacterial cell cultures adjusted to approximately 1 × 10⁶ colony forming units (CFU)/mL. Gram-positive isolates were grown in HI broth, and Gram-negative isolates in Luria–Bertani (LB) broth. Plates were incubated at 37 °C with gentle agitation (50 rpm), with the medium carefully removed via gentle pipetting and replaced with fresh sterile medium every 24 h.

### 2.4. Biofilm Characterisation

#### 2.4.1. Biofilm Enumeration

Biofilm-associated viable cells were quantified by serial dilution and plate enumeration. As per Vyas *et al* (2021)^41^, biofilms were gently washed once with sterile 1× phosphate-buffered saline (PBS), then resuspended in fresh PBS by vigorous scraping of the well surfaces. The resulting suspensions were 10-fold serially diluted in PBS, and 10 µL aliquots were spot-plated onto LB agar. Plates were incubated overnight at 37 °C, and CFU/mL of culturable viable cells were determined using Equation 1, where *df* is the dilution factor and V_d_ is the droplet volume.

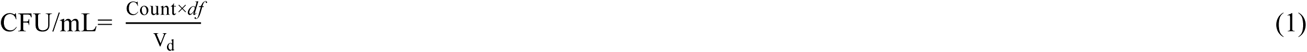

#### 2.4.2. Biofilm Biomass

Total biofilm biomass was quantified by crystal violet (CV) staining as described by Vyas *et al* (2020)^42^. Biofilms were air-dried for 30 min (or until completely dry) and fixed with 99 % methanol for 15 min. Once fixed, wells were further air-dried and stained with 0.2 % (w/v) CV (Sigma-Aldrich, USA) containing 1.9 % (v/v) ethanol for 10 min at room temperature. Excess stain was removed, and wells were gently washed twice with PBS. Bound dye was resolubilised with 1 % (w/v) sodium dodecyl sulfate (SDS) for 10 min at room temperature. Wells containing only media served as sterility controls and blanks to subtract from sample readings. The released CV dye was diluted 1:5 in 1 % SDS, and absorbance was measured at 540 nm using a CLARIOstar^®^ Plus Microplate Reader (BMG Labtech, Germany).

#### 2.4.3. Biofilm EPS

To characterise the EPS of each of these biofilms, both total EPS and major EPS components (polysaccharides, eDNA, and proteins) were investigated. Total EPS was measured using a 1,9-dimethyl-methylene blue (DMMB) dye-binding assay adapted from Peeters *et al* (2008)^43^. EPS associated polysaccharides, eDNA, and proteins were fluorescently stained according to methods outlined by Vyas *et al* (2020)^44^ and Xia *et al* (2025)^22^ using Concanavalin A conjugated to Alexa Fluor 647 (Con A–AF647; C21421, Invitrogen, USA), Sytox Blue (Molecular Probes, Invitrogen, USA), and FilmTracer SYPRO Ruby biofilm matrix stain (Molecular Probes, Invitrogen, USA), respectively. Biofilms were stained individually for 30 min in the dark with 5 µg/mL Con A–AF647, 5 µM Sytox Blue, or 0.5 × SYPRO Ruby in PBS. PBS served as a negative control to account for background fluorescence and for blank subtraction. Fluorescence was measured using a CLARIOstar^®^ Plus Microplate Reader with a 6 × 6 matrix scan to capture the non-homogeneous nature of biofilm formation on a well surface. Excitation/emission settings were as follows: Con A–AF647 (647 – 15 nm/687 – 15 nm), Sytox Blue (440 – 15 nm/484 – 20 nm), and SYPRO Ruby (450 – 15 nm/610 – 20 nm).

### 2.5. Plasma-Activated Water Generation & Biofilm Treatment

PAW was generated using a modified plasma unit from Plasmatreat (Plasmatreat GmbH, Germany). Briefly, 1.5 L autoclave sterilised deionised water was treated for 45 min in a 2 L glass Schott bottle. The plasma generation was set to a fixed power supply with variable voltage to ensure target power levels were reached for the dielectric barrier discharge at 80 watts (W) and 300 W for the gliding arc. Air was the input gas source at a flow rate of 8 and 10 litres per min (L/min), respectively (Fig. 1). Biofilms formed *in vitro* were challenged with PAW or the water control (i.e., water without plasma discharge) either immediately after PAW generation, 1-week, or 4-weeks post-generation (stored PAW/water control was placed in the dark at 4 °C and aliquots required for experimental use returned to room temperature prior to biofilm treatment). To determine the anti-biofilm efficacy of PAW, the log_10_ reduction of bacterial cells following PAW treatment was then calculated per Equation 2.

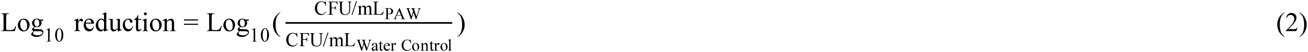

**Figure 1.**
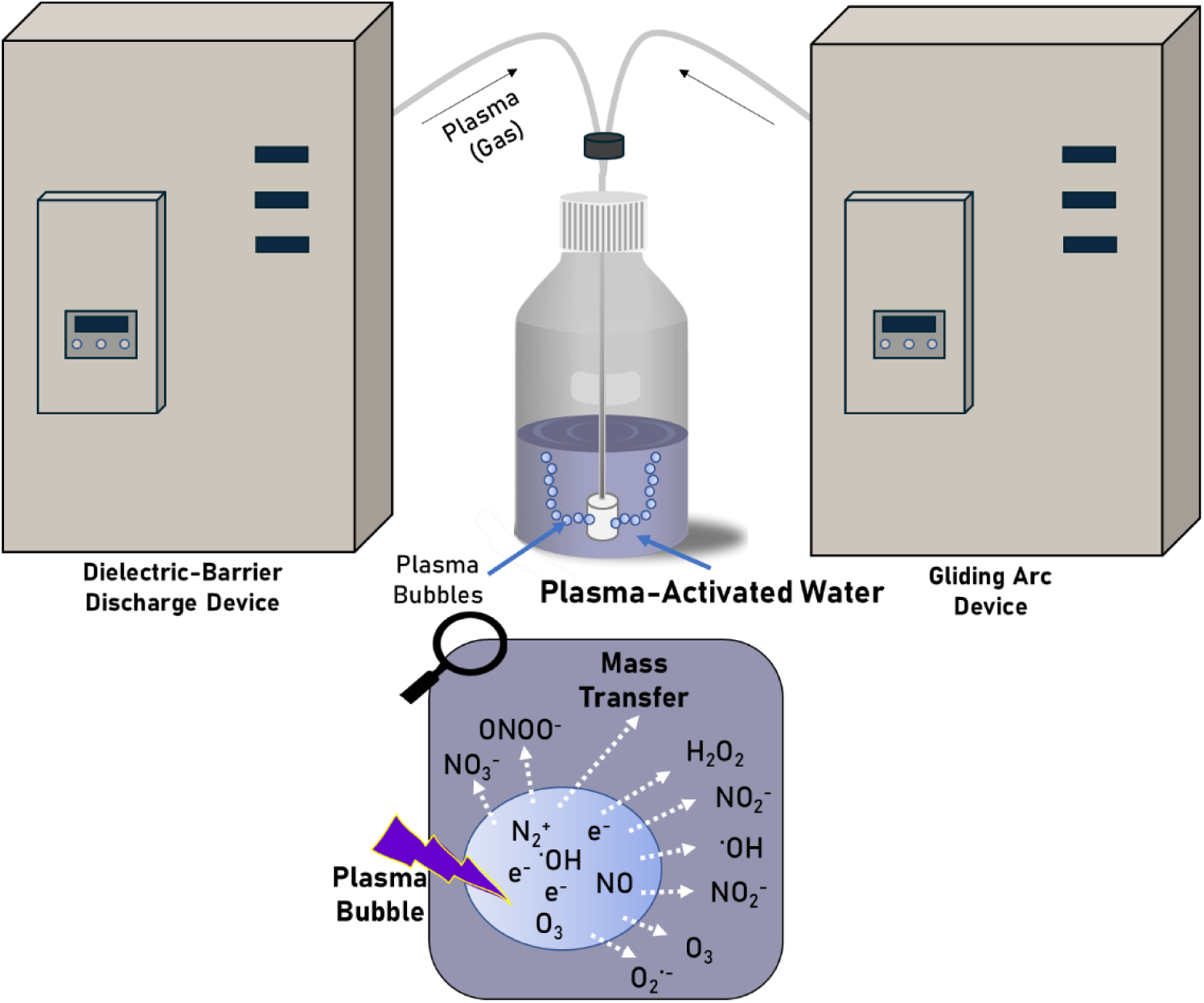
Schematic representation of the setup used in this study to generate PAW for the *in vitro* treatment of biofilms. 1.5 L of PAW was produced for 45 min using a dual-system plasma configuration comprising a dielectric barrier discharge device (80 W, air supplied at 8 L/min) and a gliding arc device (300 W, air supplied at 10 L/min). During PAW production, plasma bubbles rich in reactive species form within the water, enabling the mass transfer of these species from the gas phase (bubbles) to the liquid phase (water).

### 2.6. PAW Physicochemical Analysis

The physicochemical properties of PAW and water control such as pH, oxidation-reduction potential (ORP), and electrical conductivity were measured at ambient temperature (22-24 °C) using dedicated probes acquired from Mettler Toledo (Australia). Nitrate and nitrous acid concentration were quantified by UV-Vis spectrophotometry through application of the Beer-Lambert Law (Equation 3).

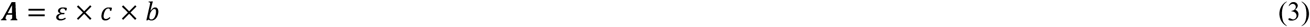

Measurements were performed using quartz cuvettes with a standard 1 cm pathlength (b). Nitrate concentration was determined from absorbance at 302 nm, and nitrous acid concentration from absorbance at 355 nm, using molar absorption coefficients (ε) of 7.24 and 23.3 L·mol⁻¹·cm⁻¹, respectively^45^. Ozone concentration was measured using a colorimetric ozone indicator assay kit (Hanna Instruments, Australia). Briefly, the control vial was filled with deionised water, while 5 mL of PAW sample was mixed with the reactive agent and 5 mL of deionised water. After 2 min, the resulting pink/reddish colour was compared with the reference colour wheel to ascertain the ozone concentration in the sample.

### 2.7. Quantification of Biofilm Reactive Oxygen Species

As per Vyas *et al* (2023)^23^, the accumulation of intracellular reactive oxygen species (ROS) was quantified using 2′,7′–dichlorofluorescin diacetate (DCFDA; Sigma-Aldrich, Australia). Briefly, biofilms were exposed for 1 min to freshly generated PAW or the water control. Following treatment, biofilms were incubated with 20 µM DCFDA (or PBS for background control) for 30 min at room temperature. Fluorescence intensity was then measured using a CLARIOstar^®^ Plus Microplate Reader for DCFDA detection (excitation/emission: 485 – 15 nm / 535 – 15 nm).

### 2.8. Scanning Electron Microscopy

Scanning electron microscopy (SEM) was utilised to examine the morphological alterations in biofilms and residing cells of *L. fusiformis*, *P. megaterium*, *E. coli*, and *P. stutzeri* following 1 min treatment with fresh PAW, relative to water control. Briefly, biofilms were grown for 48 h on 13 mm plastic Nunc Thermanox coverslips (Proscitech, USA) in a 12-well polystyrene plate, then treated with PAW or water control. Biofilms were prepared for SEM imaging using methods adapted from Vyas *et al* (2023)^23^ with the following modifications. Biofilms were pre-fixed for 30 min at 4 °C, followed by fixation for 1 h at 4 °C. After fixation, samples were washed and dehydrated through a graded ethanol series (30 %, 50 %, 70 %, and 3 × 100 %). Dehydrated biofilms were then treated with a 1:1 mixture of 100 % ethanol and hexamethyldisilazane (HMDS), followed by two successive changes of 100 % HMDS. After 30 min, the HMDS was removed and samples were allowed to air-dry. Dried biofilms were sputter-coated with a 10 nm platinum layer using a high-resolution sputter coater (Cressington Scientific Instruments, UK; 208HR) and imaged using a SU8600 cold field-emission ultra-high-resolution scanning electron microscope (Hitachi High-Tech, Japan). Images were acquired at random positions across each sample by a microscope technician blinded to the experimental conditions to minimise bias.

### 2.9. Statistical Analysis

All statistical analysis was performed using GraphPad Prism 10 (GraphPad Software, USA). Data are presented as the mean ± standard deviation (SD) or standard error of the mean (SE) as indicated. Log _10_ reduction of bacteria following PAW exposure of different treatment times and different PAW ages were compared by mixed-effects model with Šídák’s multiple comparisons test between all treatment groups, with a family-wise α threshold of 0.05. Residuals plots and QQ plots were used to assess assumptions of homoscedasticity and normality of the data, respectively. Intracellular ROS accumulation in PAW- and water control-treated biofilms were analysed by one-way ANOVA with Tukey’s post hoc test.

## 3. Results & Discussion

The bacterial isolates recovered from eggshell surfaces likely reflect a combination of poultry-derived faecal contamination, soil, and water. Identified taxa included *Staphylococcus* spp. (*S. cohnii, S. saprophyticus, S. arlettae*), *Pseudomonas* spp. (*P. putida, P. stutzeri, P. fulva*), *Escherichia coli, Acinetobacter lwoffii, Lysinibacillus fusiformis, Peribacillus frigoritolerans,* and *Priestia megaterium*. While *E. coli* is commonly a commensal of the poultry gastrointestinal tract, certain strains are associated with food safety risks, and *Salmonella* remains the primary cause of egg-associated foodborne illness^46^. Although *Salmonella* was not isolated in this study, the presence of both opportunistic pathogens and environmentally associated bacteria highlights contamination pathways relevant to egg production and handling environments warranting investigation.

### 3.1. Biofilm Characterisation of Bacteria Isolated from the Eggshell Surface

Biofilm formation is a recognised virulence and persistence mechanism among many Gram-positive and Gram-negative bacteria, including food pathogens *L. monocytogenes*, *Salmonella enterica*, *E. coli*, and thermophilic *Campylobacter*^11, 47, 48^. In food systems, biofilms pose a significant safety concern by promoting pathogen survival, cross-contamination, AMR, and overall decreased susceptibility to sanitising processes. This study examined eight bacterial isolates retrieved from the eggshell surface: four Gram-positive species (*S. arlettae*, *L. fusiformis*, *P. megaterium*, and *P. frigoritolerans*) and four Gram-negative species (*A. lwoffi*, *E. coli*, *P. fulva*, and *P. stutzeri*). Biofilm development was monitored over 24, 48, and 72 h, evaluating cultured viable cell populations, biomass, and EPS production.

All isolates formed biofilms, which varied between strains (Fig. 2). *E. coli*, a well-known and prolific biofilm former in the context of food and other diverse environments, served as a useful reference point ^48–50^. To our knowledge, the biofilm-forming abilities of several of the other species identified in this study have not been previously characterised in detail. Despite all isolates being inoculated at the same starting concentration, substantial variation in biofilm cell density was observed. For instance, *P. megaterium* maintained relatively low viable populations (≈ 6.5 log CFU/mL), while *E. coli* reached and sustained > 8.2 log CFU/mL (Fig. 2A), underscoring species-specific differences in biofilm growth and maturation dynamics. Some strains, such as *P. megaterium* and *P. fulva*, maintained stable cell densities throughout the biofilm maturation process, while others, including *A. lwoffi* and *L. fusiformis*, showed a decline at 72 h. Biofilm development typically progresses through four key stages: (i) reversible attachment of planktonic cells, (ii) irreversible attachment accompanied by EPS production, forming the scaffold for initial three-dimensional structure development, (iii) maturation, characterised by phenotypic and genetic diversification that enhances structural resilience against biological, physical, and chemical stressors, and (iv) dispersal, where cells detach and return to the planktonic state^51, 52^. The variation in viable cell populations observed among isolates over time likely reflects both inter-genus (e.g., *S. arlettae* vs *P. fulva*) and intra-genus (e.g., *P. fulva* vs *P. stutzeri*) differences in biofilm development, driven by distinct physiological traits, metabolic strategies, and regulatory mechanisms. These intrinsic factors interact with fluctuating local microenvironmental variables such as pH, temperature, nutrient and gas availability, as well as waste and quorum sensing molecule accumulation, collectively shaping how efficiently and rapidly each strain transitions through the biofilm life cycle and oscillates between sessile biofilms and dispersed, free, single planktonic cell states.

**Figure 2.**
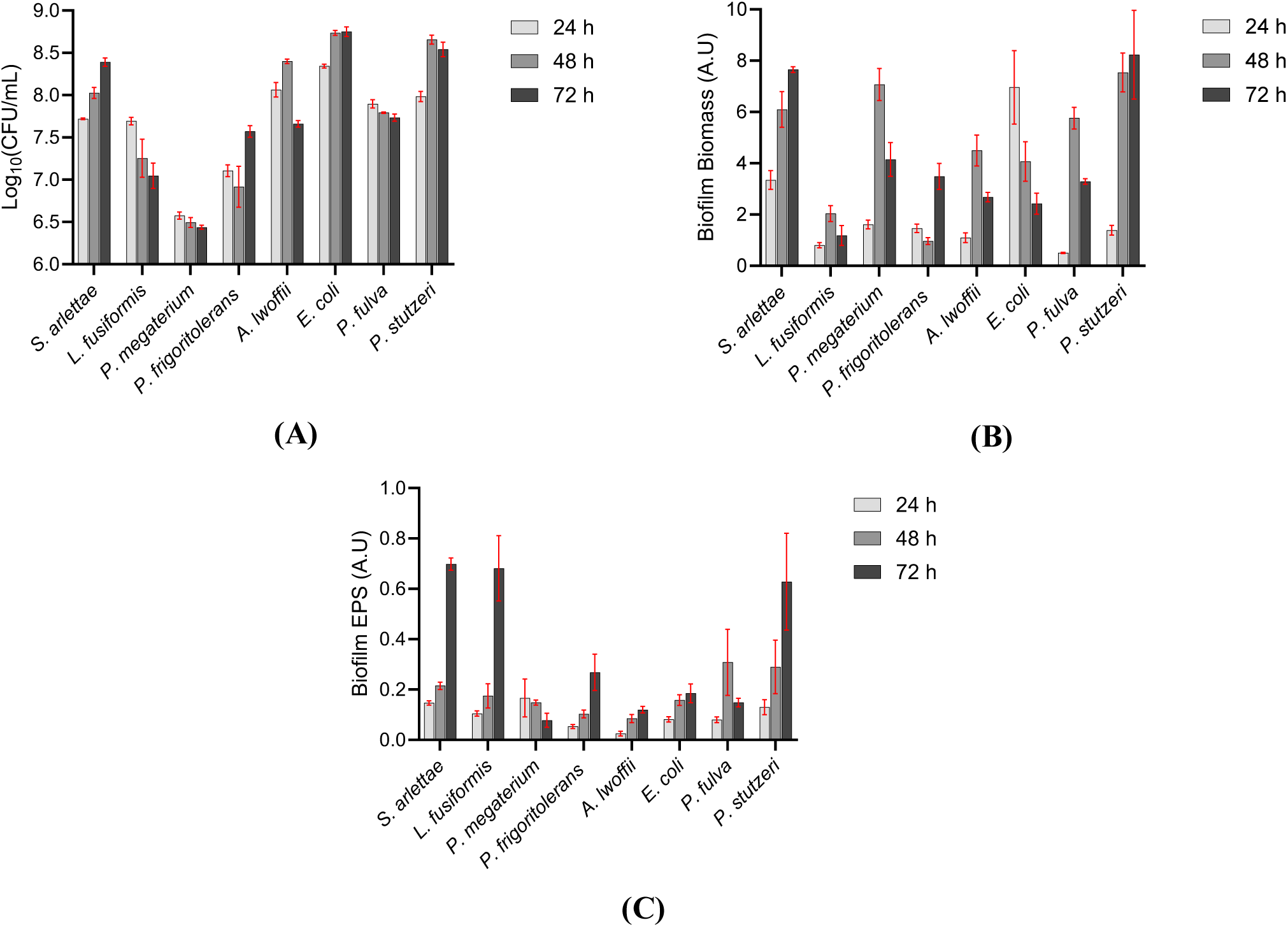
Biofilms formed at 24, 48, and 72 h by eight bacterial isolates from eggshell surfaces were characterised for biofilm growth, biomass, and EPS. **(A)** Biofilm growth was assessed by viable cell enumeration of culturable cells, **(B)** biofilm biomass was quantified using CV staining, and **(C)** biofilm EPS was measured using DMMB staining. Data are presented as mean ± SE; *n* = 3 biological replicates with 2 technical replicates each.

To further investigate differences in biofilm development, total biofilm biomass was quantified using CV staining. Biomass measurements mirrored culturable viable cell trends for *S. arlettae* (Fig. 2B), suggesting that the enumerated live cells were the primary contributors to the total biomass detected. In contrast, other strains exhibited disproportionately higher biomass relative to viable cell counts, indicating greater contributions from EPS and/or viable but non-culturable and non-viable cells. Given that the EPS is documented to constitute up to 80–85% of total biofilm biomass and serves as a key protective barrier against physical (e.g., shear stress, desiccation), biological (e.g., host immune-type stresses, regulatory disruption), and chemical (e.g., enzymatic degradation, antimicrobial agents) challenges^53, 54^, its production was further quantified using DMMB staining. EPS matrix production was confirmed in all isolates (Fig. 2C), generally increasing with biofilm age. Overall, these findings demonstrate that biofilm formation among eggshell-associated bacteria is isolate-dependent, with the EPS matrix likely playing a central role in biofilm architecture and stability.

The EPS matrix comprises a complex and dynamic “matrixome” that defines both its compositional and functional diversity^54, 55^. EPS is primarily composed of polysaccharides, eDNA, proteins, and lipids. As all isolates in this study produced an EPS matrix and given the limited knowledge of these bacterial biofilms, we sought to characterise EPS composition and determine whether biofilm age influenced the abundance and presence of each component. Across bacterial biofilms, polysaccharides typically represent the most abundant EPS constituent, forming the bulk of the matrix and contribute to adhesion, scaffolding, and structural stability, protecting embedded microbial cells^54–56^. Polysaccharide production remained relatively consistent among strains over the assessed period of biofilm growth; however, *S. arlettae* and *P. megaterium* displayed peak polysaccharide abundance at 24 h, followed by a moderate decline and stabilisation between 48–72 h (Fig. 3A). eDNA followed a similar trend (Fig. 3B), consistent with its recognised structural role in biofilm cohesion and scaffolding in relatively better-characterised biofilms of bacteria like *P. aeruginosa* and *E. coli*^57, 58^. Beyond its structural function, eDNA can serve as a nutrient source and facilitate horizontal gene transfer, thereby enhancing genetic diversity and potentially contributing to AMR^55, 57^. Protein content increased markedly in all strains at 48–72 h (Fig. 3C), suggesting that proteins become increasingly important during biofilm maturation. Like polysaccharides and eDNA, proteins support cell–cell and surface adhesion and can enhance protection against antimicrobial agents^59, 60^. Lipids were not examined in this study, but their roles in structural integrity, stress tolerance, and reducing the biofilms antimicrobial susceptibility warrant further investigation^55^.

**Figure 3.**
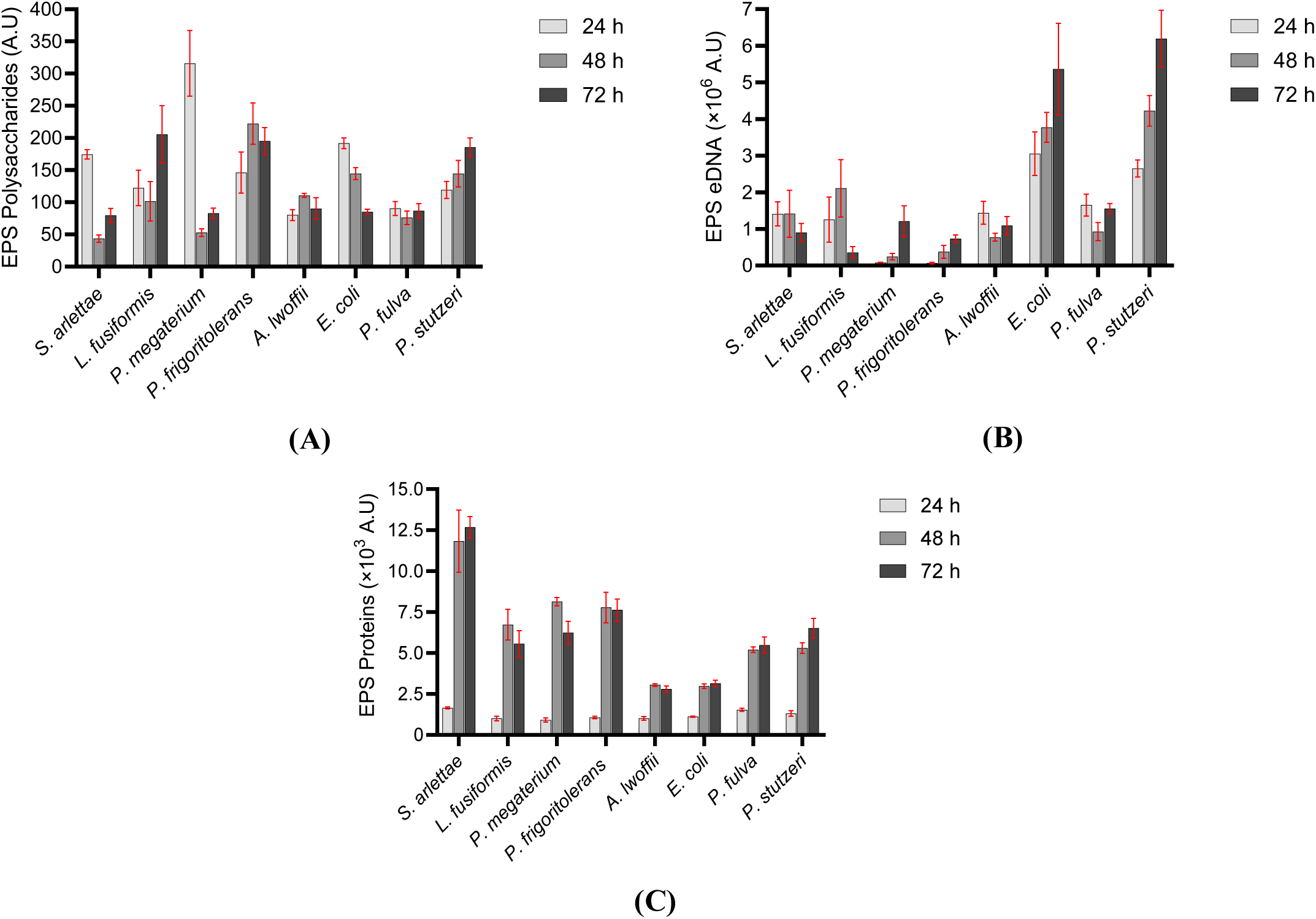
Profiling of EPS composition within biofilms formed at 24, 48, and 72 h by eight bacterial isolates from eggshell surfaces revealed the presence of matrix-associated polysaccharides, eDNA, and proteins. Fluorescent staining was used to quantify component-specific fluorescence signals: **(A)** Con A–AF647 for EPS-associated polysaccharides, **(B)** Sytox Blue for eDNA, and **(C)** SYPRO Ruby for proteins. Data are presented as mean ± SE; *n* = 3 biological replicates with 2 technical replicates each.

Because EPS components do not accumulate uniformly, fluctuating with biofilm age, we examined the relative contributions of polysaccharides, eDNA, and proteins individually over time. Such age-dependent shifts in matrix composition are likely to influence biofilm susceptibility to antimicrobials ^61^, thereby potentially reducing the efficacy of conventional and emerging, novel antimicrobials and stringent cleaning and decontamination procedures. Targeting specific EPS components has already proven effective; for example, DNase treatment disrupts early-stage biofilms of *P. aeruginosa*, *S. aureus*, and *L. monocytogenes*, demonstrating that matrix constituents themselves represent viable intervention targets in foo d environments^62–64^. Moreover, EPS-rich biofilms have been repeatedly associated with decreased efficacy of standard food-industry sanitisers, underscoring the importance of matrix profiling which may inform the timing and selection of cleaning and disinfection regimes and agents^60^. Collectively, these findings emphasise that understanding both EPS composition and temporal dynamics is informative when developing effective sanitisers and cleaning strategies to mitigate biofilm persistence in food production systems.

### 3.2. Evaluation of the Anti-Biofilm Efficacy of PAW

PAW is well established for its potent antimicrobial activity against planktonic bacteria across sectors including medicine, water treatment, and agriculture^21–25, 35, 65–67^. Within the food industry, PAW and other CAP-related technologies (e.g., plasma jets and pens) have emerged as promising sanitising tools that enhance food safety, purportedly without leaving harsh, persistent chemical residues typical of conventional disinfecting agents such as bleach, while maintaining product quality^20, 25, 27, 68^. Several studies have demonstrated the rapid bactericidal activity of PAW against planktonic foodborne pathogens such as *E. coli*, *S. enterica*, *Salmonella* Typhimurium, and *L. monocytogenes*, which are commonly associated with outbreaks and recalls involving fresh produce, poultry, meat, and food-contact surfaces^25, 27, 33, 35^. Recent research has shifted toward balancing antimicrobial efficacy with operational and economic feasibility to optimise PAW for industrial implementation. Investigations have examined altering the input gas composition (e.g., pure oxygen versus air) when generating PAW to finetune produced RONS, activation medium (e.g., tap versus deionised water), and application parameters such as PAW treatment mode (e.g., dipping, spraying, washing) and treatment duration (seconds to hours) for large-scale processing^24, 25, 33^. Despite this progress, few studies have assessed the anti-biofilm efficacy of PAW in egg decontamination within the poultry industry, even though biofilms impose a global economic burden estimated at approximately US$324 billion annually in the food sector^69^.

In this study, we evaluated the effects of short PAW exposure times (1, 5, and 15 min) and PAW age (fresh, 1-week-old, and 4-week-old) on 48 h biofilms formed by both Gram-positive (*L. fusiformis*, *P. megaterium*) and Gram-negative (*E. coli*, *P. stutzeri*) isolates. A 48 h biofilm was selected as the standard condition for antimicrobial susceptibility testing as it produced consistent and mature biofilms based on our viable cell counts, biomass, and EPS measurements. PAW effectively inactivated bacteria in biofilms formed by all four species to varying degrees depending on PAW age and exposure time (Fig. 4). Relative to the water control, reductions in Gram-positive biofilms were significant but moderate (≈ 3–4.5 log CFU/mL), even with freshly generated PAW and extended exposure (15 min; Fig. 4A and B). In contrast, fresh PAW rapidly and significantly reduced Gram-negative *E. coli* and *P. stutzeri* biofilm viability within 1 min (≈ 7.5–8 log CFU/mL) below the limit of detection (Fig. 4C and D).

**Figure 4.**
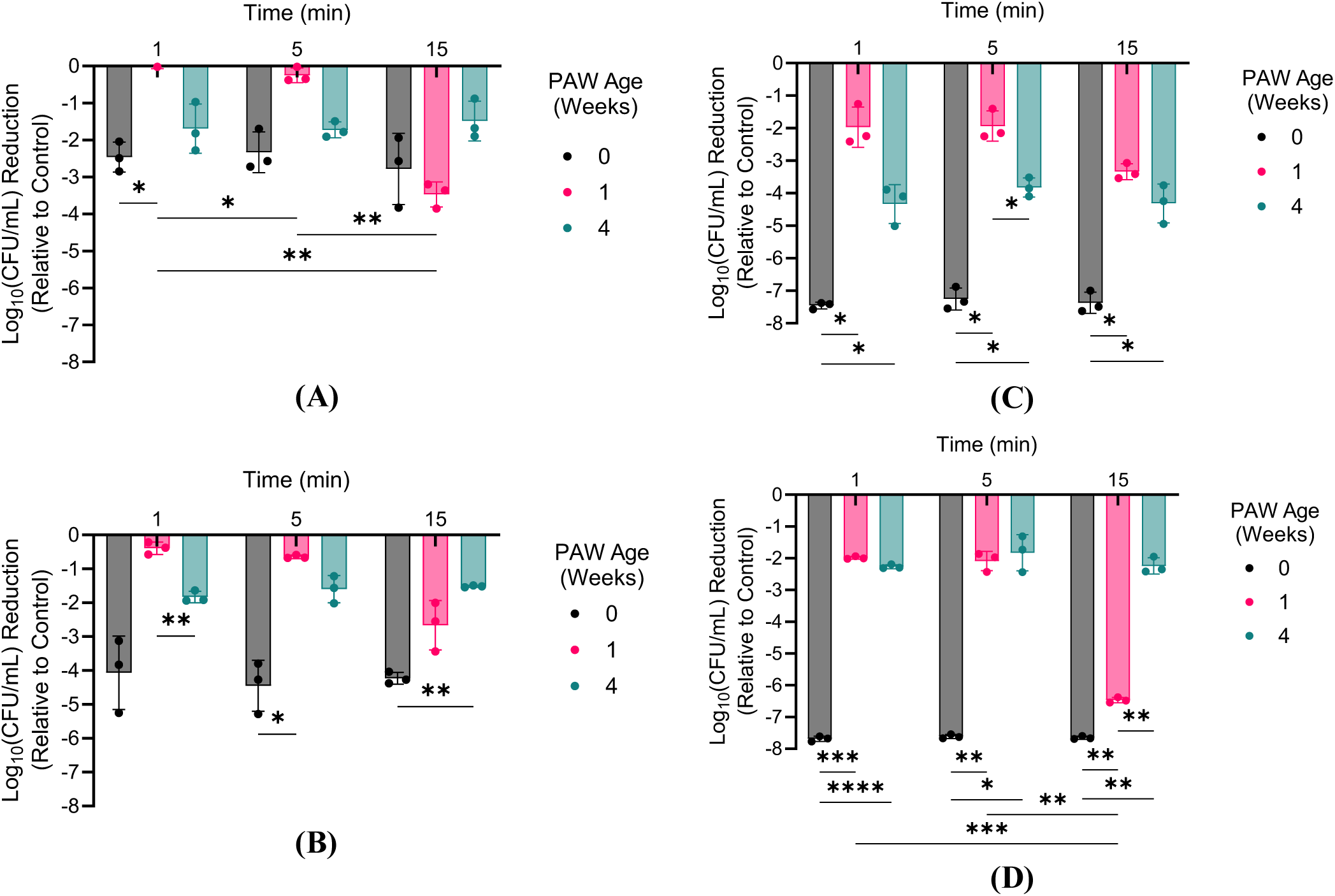
PAW treatment was most effective against biofilms of the Gram-negative bacteria *E. coli* and *P. stutzeri*. 48 h biofilms of (A) *L. fusiformis*, (B) *P. megaterium*, (C) *E. coli*, and (D) *P. stutzeri* were exposed for 1, 5, and 15 min to freshly generated PAW (0 weeks old) or PAW aged for 1- and 4-weeks. The depicted reduction in biofilm viability was determined relative to biofilms treated with the respective water controls. Data are presented as mean ± SD; \**P* ≤ 0.05, \*\**P* ≤ 0.01, \*\*\**P* ≤ 0.001, and \*\*\*\**P* ≤ 0.0001; *n* = 3 biological replicates with 2 technical replicates each.

Differences in PAW susceptibility between Gram classifications have been attributed to variations in cell envelope architecture. Gram-positive bacteria possess a thick (20–80 nm) peptidoglycan layer that provides strong mechanical and chemical resistance, whereas Gram-negative species have a thinner (< 10 nm) wall but an additional outer membrane with porins and surface appendages that can facilitate the penetration of reactive species. These structural distinctions influence tolerance to environmental stressors such as heat, ultraviolet radiation, and the RONS generated by plasma treatment. For instance, Mai-Prochnow *et al* (2016)^70^ reported that biofilms of the Gram-positive *Bacillus subtilis* (55 nm cell wall) were highly resistant to CAP delivered via the kINPen device, showing only a 1 log_₁₀_ reduction after 10 min, whereas *P. aeruginosa* (2.4 nm cell wall) biofilms were completely eradicated under identical conditions. In multi-species biofilms of *P. aeruginosa* and *Staphylococcus epidermidis*, the latter, a Gram-positive species, also resisted CAP-induced hydroxyl and oxygen radical damage to the cell wall. More recently, Xia *et al* (2025)^22^ demonstrated that *in situ* PAW treatment, where biofilms formed on stainless-steel coupons, mimicking food processing surface contamination, were exposed to PAW directly during plasma activation, resulting in substantial inactivation of both *E. coli* and *S. aureus* biofilms, with *S. aureus* inactivated more rapidly. These findings suggest that plasma delivery mode (e.g., PAW versus plasma plume) and treatment interface strongly influence biofilm susceptibility. The comparatively greater tolerance of the Gram-positive biofilms observed in this study likely reflects the reduced reactivity of PAW even when applied immediately post-generation, rather than *in situ* during plasma activation. Many short-lived reactive species that contribute to the antimicrobial efficacy of PAW, with lifetimes ranging from nanoseconds to seconds, are rapidly lost under these conditions ^21, 24^. This loss likely also explains the diminished efficacy of aged PAW (1- and 4-week-old samples), as the decay of transient RONS over time reduces antimicrobial potency; consequently, extending treatment beyond 5 min conferred no additional benefit as seen for some of the PAW-treated biofilms.

Given the demonstrated antimicrobial and anti-biofilm efficacy of PAW, its physicochemical properties were analysed immediately after generation (fresh PAW) and following 1- and 4-weeks of storage to identify any associations with bactericidal activity. While both short- and long-lived RONS, such as ozone, nitrite, and nitrate, are widely recognised as primary contributors to PAW-mediated antimicrobial effects, plasma activation also induces broader physicochemical changes, including decreases in pH with concurrent increases in ORP and conductivity (Table 1).

**Table 1.**
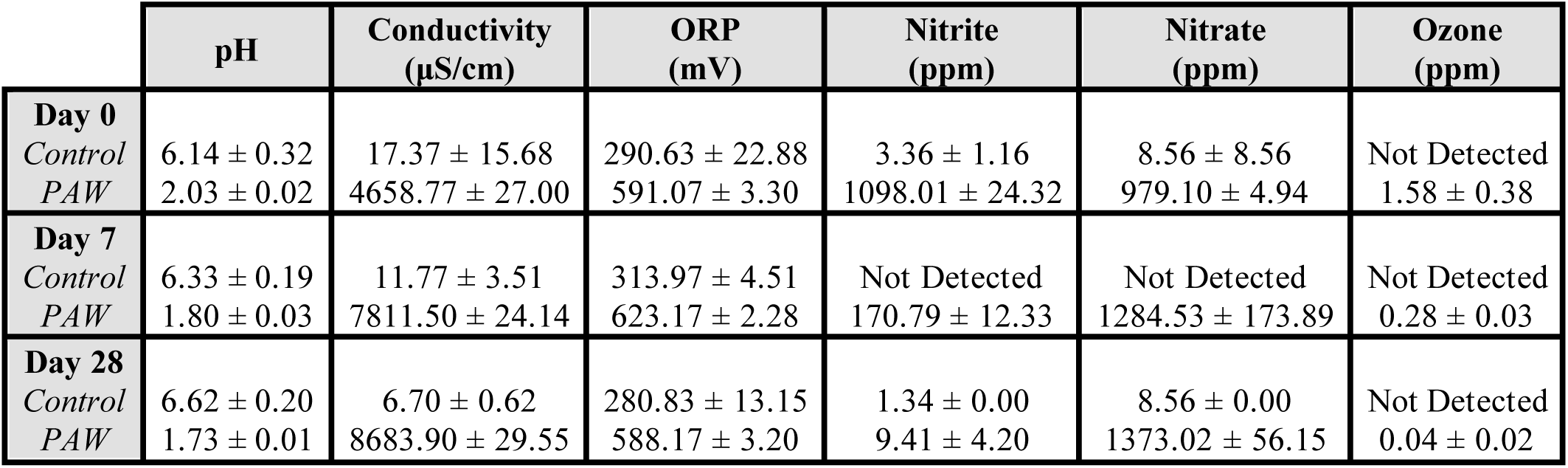
Physicochemical properties and RONS concentrations of PAW measured immediately after generation (fresh) and at 7 days (1-week-old) and 28 days (1-week-old) of storage. A water control is included for comparison. Values represent mean ± SD from 3 independent measurements.

| | pH | Conductivity<br>(μS/cm) | ORP<br>(mV) | Nitrite<br>(ppm) | Nitrate<br>(ppm) | Ozone<br>(ppm) |
| --- | --- | --- | --- | --- | --- | --- |
| **Day 0** |  |  |  |  |  |  |
| Control | $6.14 \pm 0.32$ | $17.37 \pm 15.68$ | $290.63 \pm 22.88$ | $3.36 \pm 1.16$ | $8.56 \pm 8.56$ | Not Detected |
| PAW | $2.03 \pm 0.02$ | $4658.77 \pm 27.00$ | $591.07 \pm 3.30$ | $1098.01 \pm 24.32$ | $979.10 \pm 4.94$ | $1.58 \pm 0.38$ |
| **Day 7** |  |  |  |  |  |  |
| Control | $6.33 \pm 0.19$ | $11.77 \pm 3.51$ | $313.97 \pm 4.51$ | Not Detected | Not Detected | Not Detected |
| PAW | $1.80 \pm 0.03$ | $7811.50 \pm 24.14$ | $623.17 \pm 2.28$ | $170.79 \pm 12.33$ | $1284.53 \pm 173.89$ | $0.28 \pm 0.03$ |
| **Day 28** |  |  |  |  |  |  |
| Control | $6.62 \pm 0.20$ | $6.70 \pm 0.62$ | $280.83 \pm 13.15$ | $1.34 \pm 0.00$ | $8.56 \pm 0.00$ | Not Detected |
| PAW | $1.73 \pm 0.01$ | $8683.90 \pm 29.55$ | $588.17 \pm 3.20$ | $9.41 \pm 4.20$ | $1373.02 \pm 56.15$ | $0.04 \pm 0.02$ |

Freshly generated PAW exhibited a strongly acidic pH (pH 2.03 ± 0.02, 22.7 ± 0.1 °C) compared to the water control (pH 6.1 ± 0.3, 23.1 ± 0.3 °C), indicating that plasma activation alters the pH without significant increases in temperature under the applied generation conditions. PAW remained highly acidic throughout storage, with a modest decrease in pH observed over time (pH 2.03 initially to pH 1.73 after 4-weeks). While acidic conditions of the PAW may contribute additional stress to bacterial cells, pH alone is unlikely to be the primary driver of the anti-biofilm activity demonstrated. Supporting this interpretation, Vyas *et al* (2023)^23^ reported that non-plasma-treated water adjusted to the same acidic pH as PAW (pH 2.8) did not significantly reduce *E. coli* biofilm viability, suggesting that RONS detected in PAW were central to antimicrobial efficacy. Future studies utilising pH-matched controls would further clarify the contribution of acidity to antimicrobial outcomes observed here.

Fresh PAW also demonstrated markedly higher electrical conductivity (4658.8 ± 27.0 µS/cm) and ORP (591.1 ± 3.3 mV) relative to the water control (17.4 ± 15.7 μS/cm and 290.6 ± 22.9 mV, respectively). Conductivity increased progressively during storage, nearly doubling after 4-weeks (8683.90 ± 29.55 μS/cm), whereas ORP remained relatively stable over time. Notably, increased conductivity did not correspond to enhanced antimicrobial activity, as aged PAW generally exhibited reduced bactericidal efficacy, indicating that conductivity alone is not predictive of antimicrobial potency.

Consistent with previous literature, RONS including ozone, nitrate, and nitrite were detected in the PAW generated for this study, while concentrations in the water control were negligible or below detection limits. The highest RONS concentrations were observed immediately after PAW generation (in ppm; ozone: 1.58 ± 0.38; nitrate: 979.10 ± 4.94; nitrite: 1098.01 ± 24.32). In contrast, ozone and nitrite concentrations declined during storage, consistent with the reduced antimicrobial efficacy of aged PAW. While several of the RONS detected in PAW, including ozone and nitrate, are considered relatively longer-lived species with reported half-lives ranging from minutes to years^20^, the stability is highly dependent on environmental conditions such as pH, temperature, and solution composition^71^. Given the strongly acidic pH and highly reactive environment of PAW as determined upon physicochemical analysis, ozone likely decayed rapidly following generation. Precise decay kinetics could not be resolved for any species within these time scales, so future studies monitoring ozone, nitrite, and nitrate concentrations over shorter timeframes (< 1 h to within days post-generation) would provide greater insight into their stability and provide additional understanding into the anti-biofilm activity of PAW. In contrast to ozone and nitrite, which decline during storage, nitrate concentrations increased, reflecting the oxidative conversion of nitrite to nitrate, facilitating the continued acidification of aged PAW.

Collectively, these observations support PAW-associated RONS, rather than bulk physicochemical parameters alone, as the primary drivers of bactericidal and anti-biofilm activity. To more precisely delineate the contribution of individual reactive species to microbial inactivation, future work incorporating selective scavenger assays would be valuable. For instance, ozone can be selectively scavenged using uric acid added to the water prior to plasma activation^24^. However, as ozone can generate secondary reactive species, including hydroxyl radicals, hydroxide ions, molecular oxygen, and superoxide anions, additional targeted scavengers may be required. For instance, Tiron, a well-established scavenger of superoxide anions, has been successfully utilised in previous studies to interrogate PAW-mediated antimicrobial mechanisms^21, 23–25^. Notably, selective scavenging of ROS such as superoxide has been shown to significantly reduce the antimicrobial efficacy of PAW against planktonic cells and biofilms of foodborne pathogens including *E. coli, L. monocytogenes,* and *S. enterica*^21, 23–25^. In some cases, removal of key reactive species from the PAW resulted in biofilm viability indistinguishable to viability of water-treated controls, underscoring the central role of RONS in PAW-mediated antimicrobial activity^23^.

### 3.3. Mechanistic Insights into the Anti-Biofilm Activity of PAW

Physicochemical analysis confirmed the presence of ozone, nitrite, and nitrate in PAW. These species are well documented for their potent antimicrobial effects against both planktonic cells and biofilms of key foodborne pathogens such as *E. coli*, *P. aeruginosa*, *L. monocytogenes*, *S. aureus*, and *S. enterica*^22, 24, 25, 28, 33, 72^. Given the abundance and diversity of RONS detected in PAW, and the role in plasma-induced disruption of cell membranes, nucleic acids, proteins, and other intracellular targets^21–23, 72^, subsequent analyses evaluated the mechanistic contribution of these species to the antimicrobial activity of freshly-generated PAW against eggshell-associated bacteria and their biofilms.

#### 3.3.1. PAW Alters Biofilm Architecture & Individual Cell Morphology

SEM was used to qualitatively assess the effects of PAW treatment on bacterial biofilm cells and overall biofilm architecture (Fig. 5). Among the Gram-positive isolates, *L. fusiformis* (Fig. 5A) and *P. megaterium* (Fig. 5B), the predominant effect of PAW appeared to be extensive removal of the EPS matrix, as indicated by visibly reduced extracellular material compared with the water-treated controls. This is a promising finding, as the EPS often provides significant protection against external stressors, including antimicrobial agents. Future research could explore the application of PAW as a pre-treatment wash, followed by conventional industry-standard sanitisers (e.g., chlorine-based agents), to improve the removal of resilient biofilms. This approach may be particularly beneficial for Gram-positive bacteria, which demonstrated greater tolerance to PAW in this study. Supporting this approach, Vyas *et al* (2023)^23^ reported that, in biomedical applications targeting chronic wound, using PAW as a pre-treatment significantly enhanced the anti-biofilm efficacy of subsequently applied topical antiseptics, resulting in complete biofilm eradication. Notably, this level of efficacy was not achieved when either PAW or antiseptics were utilised alone, as both typically exhibit reduced activity against biofilms in chronic wound infections. In contrast, the Gram-negative biofilms of *E. coli* (Fig. 5C) and *P. stutzeri* (Fig. 5D) did not display a consistent response attributable to Gram classification. Instead, PAW induced distinct morphological alterations, *E. coli* cells exhibited pronounced surface deformation, including shrinkage and crumpling, whereas *P. stutzeri* biofilms showed localised cell detachment accompanied by partial removal of the EPS matrix.

**Figure 5.**
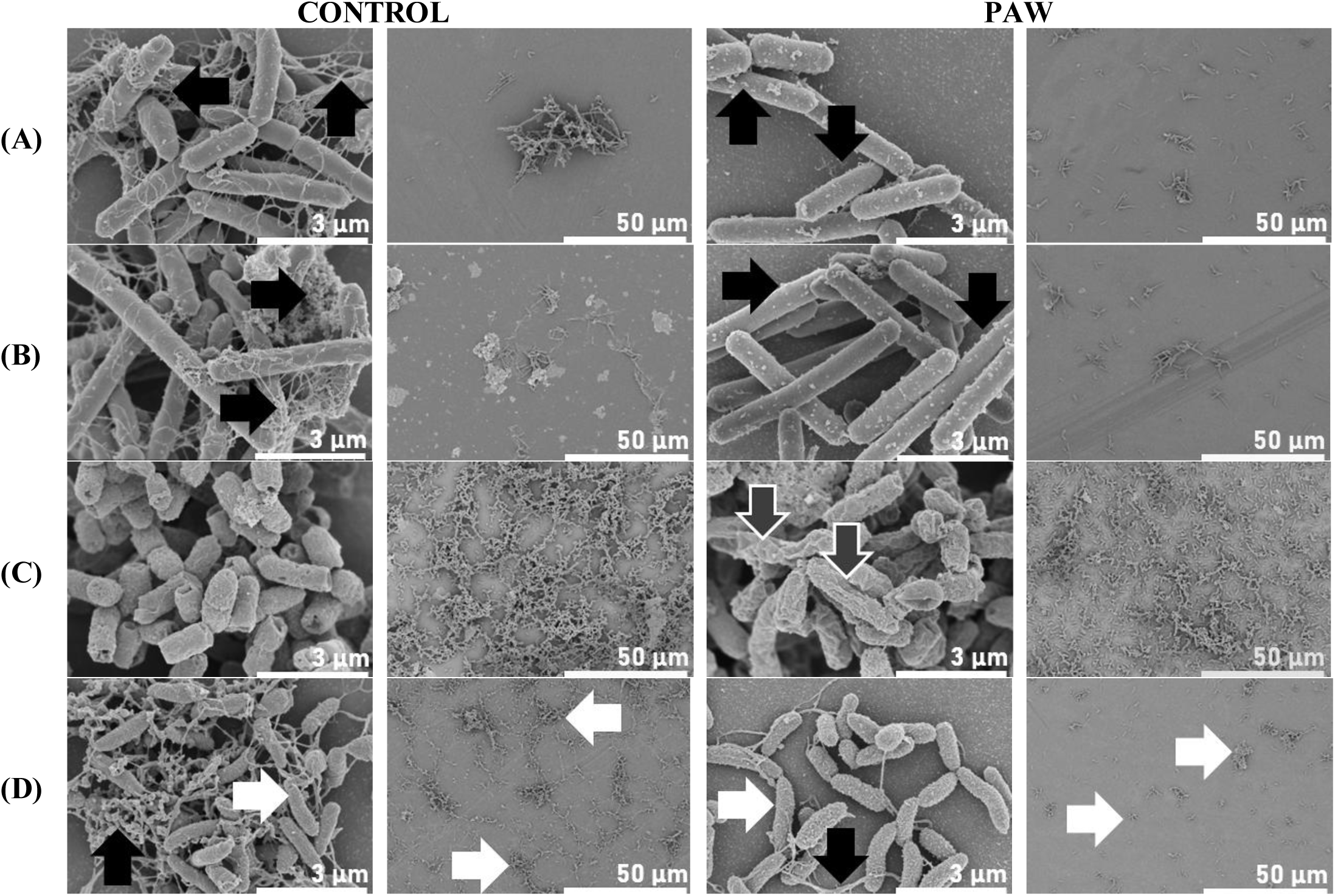
SEM reveals distinct morphological and structural alterations in bacterial biofilms and their cells following PAW treatment. 48 h biofilms of (A) *L. fusiformis*, (B) *P. megaterium*, (C) *E. coli*, and (D) *P. stutzeri* were exposed to freshly generated PAW or water control for 1 min. Observed morphological changes included EPS reduction (black arrows), cell surface shrinkage and deformation (grey arrows), and localised removal of biofilm cells (white arrows).

To our knowledge, this study is the first to visualise the effects of PAW on *L. fusiformis*, *P. megaterium*, and *P. stutzeri* biofilms, with *E. coli* serving as a useful point of reference which has been previously characterised elsewhere in the context of PAW. Vyas *et al* (2023)^23^ reported similar findings in 48 h *E. coli* biofilms, where 15 min PAW exposure caused noticeable cell damage including cell flattening and membrane blebbing indicative of outer membrane vesicle release as a bacterial survival response to oxidative stress generated. These morphological alterations were consistent with increased membrane permeability and depolarisation detected within 1 min of PAW exposure, confirming that substantial membrane damage induces rapid cell death^23^. Building on these insights, future investigations should examine PAW-induced membrane effects in our eggshell-associated bacteria, with transmission electron microscopy potentially providing additional deeper insights into both membrane integrity and intracellular alterations. Overall, pronounced physical disruption of biofilm structure and cellular morphology was observed following brief exposure to PAW, underscoring its strong anti-biofilm activity.

#### 3.3.2. Intracellular Accumulation of PAW-Associated ROS May Contribute to Lethality

PAW-derived RONS have been shown to accumulate intracellularly within treated biofilms of both Gram-positive and Gram-negative bacteria^21–24^. Xia *et al* (2025)^22^ demonstrated that PAW disrupts polysaccharide components of the biofilm matrix that are essential for structural integrity in *E. coli* and *S. aureus* biofilms, likely facilitating deeper diffusion of RONS into the biofilm and resulting in extensive membrane damage and rapid cell death. Similarly, Vyas *et al* (2025)^21^ reported intracellular RONS accumulation in PAW-treated *E. coli* biofilms is linked to the significant upregulation of 478 genes associated with antioxidant defence (e.g., thioredoxin), metabolism, and membrane transporter proteins to shuttle out RONS, reflecting bacterial survival responses under PAW-induced oxidative stress.

In the present study, we observed intracellular accumulation of ROS within *E. coli* biofilms following 1 min of PAW treatment (Fig. 6). While elevated ROS levels were also detected in *P. stutzeri* and *P. megaterium* biofilms relative to water-treated controls, these increases were not statistically significant. This aligns with the reduced susceptibility of Gram-positive biofilms (*L. fusiformis* and *P. megaterium*) to short PAW exposures. However, given the notable viability loss observed for *P. stutzeri* biofilms under the same conditions, the absence of significant intracellular ROS accumulation is intriguing. Follow-up studies could investigate whether longer PAW treatment durations enhance intracellular ROS accumulation. Moreover, previous work has shown that reactive nitrogen species (RNS) accumulate within PAW-treated biofilms; notably, Vyas *et al* (2023)^23^ reported that intracellular RNS levels exceeded ROS in *E. coli* biofilms following PAW exposure, underscoring the importance of understanding both species groups in PAW-mediated antimicrobial activity.

**Figure 6.**
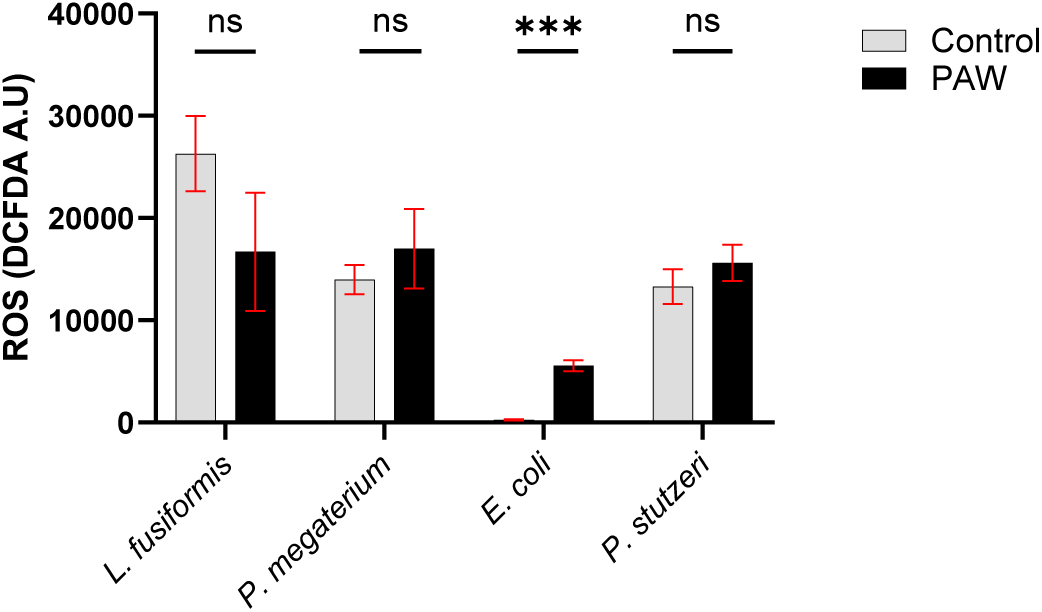
*E. coli* biofilms exhibit significant intracellular ROS accumulation following PAW treatment. Intracellular ROS levels were detected and quantified in 48 h biofilms of *L. fusiformis*, *P. megaterium*, *E. coli*, and *P. stutzeri* using DCFDA staining after 1 min treatment with freshly generated PAW, compared to water-treated controls. Data are presented as mean ± SE; \*\*\**P* ≤ 0.001 and ns *P* > 0.05; *n* = 3 biological replicates with 2 technical replicates each.

## 4. Conclusions

This study presents the first comprehensive characterisation of the biofilm-forming abilities of bacteria isolated from chicken eggshells and the utility of PAW as a decontamination strategy against these bacteria. Future investigations should examine these isolates within polymicrobial biofilms, as naturally occurring biofilms are generally considered to exist as mixed-species communities. Although *Salmonella* and *Listeria* species, major agents of egg-associated foodborne illness, were not detected in this study, their inclusion in future work remains highly relevant to the field of food safety and security. Additionally, using contextually relevant models, such as biofilms formed directly on eggshells, will improve the translational value of our PAW efficacy data, as growth and antimicrobial susceptibility most likely differ markedly for biofilms developed *in vitro* on laboratory plastics such as microtiter plates^14, 73^. Although antimicrobial efficacy declined with PAW age due to reactive species decay, optimising PAW generation and application conditions will be critical for its practical utilisation as a non-thermal sanitisation strategy in food processing environments. Further elucidation of reactive species diversity beyond those identified in this study, along with the chemical pathways governing their formation and reactions in PAW, will also advance our understanding of its translational potential. Nonetheless, this study demonstrates that PAW exhibits potent anti-biofilm activity driven by RONS, bringing PAW a step closer to potentially improving food safety, reducing food loss and waste, and decreasing our reliance on conventional industrial antimicrobials.

## Acknowledgements

This work was financially supported by Adelaide University, Australia Deputy Vice Chancellor’s Agrifood and Wine FAME Funding Strategy, the End Food Waste Cooperative Research Centre, Australia and Australia’s Economic Accelerator – Ignite grant (IG240100251). K. R. receives a Future Making Fellowship by the Division of Research and Innovation, Adelaide University, Australia. The authors gratefully acknowledge the expert technical assistance and support of Dr Nobuyuki Kawashima and Mr Christopher Leigh at Adelaide Microscopy (Adelaide University) and Microscopy Australia. We also thank Mr Connor Jessop (PhD Candidate, Adelaide University) for his invaluable technical support. We thank Mr John Byrne and Mrs Catriona Byrne of Feather & Peck Pty. Ltd. (Australia) for their generous donation of in-kind project resources.

## 5. Author Contributions

Conceptualisation, H.K.N.V.; methodology, H.K.N.V.; formal analysis, H.K.N.V., A.I.A., B.H.K.; experimental investigation, H.K.N.V., B.H.K., A.B., S.J.; data curation, H.K.N.V., A.I.A., B.H.K.; visualisation, H.K.N.V., A.I.A., B.H.K.; writing—original draft preparation, H.K.N.V. A.I.A., B.H.K., S.J.; writing—review and editing H.K.N.V., B.H.K., S.J., A.R.M., B.R.C., K.R.; supervision, K.R. and H.K.N.V.; resources, K.R., B.R.C., and A.R.M.; funding acquisition, K.R., B.R.C., and A.R.M. All authors have read and agreed to the published version of the manuscript.

